# Reduction in CD11c^+^ microglia correlates with clinical progression in chronic experimental autoimmune demyelination

**DOI:** 10.1101/396036

**Authors:** Florian Mayrhofer, Zhanna Dariychuk, Anthony Zhen, Daniel J. Daugherty, Peter Bannerman, Angela M. Hanson, David Pleasure, Athena Soulika, Wenbin Deng, Olga V. Chechneva

## Abstract

Multiple sclerosis (MS) is a chronic autoimmune demyelinating disease with high variability of clinical symptoms. In most cases MS appears as a relapsing-remitting disease course that at a later stage transitions into irreversible progressive decline of neurologic function. The mechanisms underlying MS progression remain poorly understood. Experimental autoimmune encephalomyelitis (EAE) is an animal model of MS. Here we demonstrate that mice that develop mild EAE after immunization with myelin oligodendrocyte glycoprotein 35-55 are prone to undergo clinical progression around 30 days after EAE induction. EAE progression was associated with reduction in CD11c^+^ microglia and dispersed coalescent parenchymal infiltration. We found sex-dependent differences mediated by p38α signaling, a key regulator of inflammation. Selective reduction of CD11c^+^ microglia in female mice with CD11c-promoter driven *p38α* knockout (*KO)* correlated with increased rate of EAE progression. In protected animals, we found CD11c^+^ microglia forming contacts with astrocyte processes at the glia limitans and immune cells retained within perivascular spaces. Together, our study provides evidence on the protective role of CD11c^+^ microglia in controlling CNS immune cell parenchymal infiltration in autoimmune demyelination.

## Introduction

Multiple sclerosis (MS) is an autoimmune demyelinating disorder characterized by myelin loss, chronic inflammation, and axonal degeneration. MS is a complex disease with variable outcomes. In 80% of patients, MS appears as a relapsing-remitting form (RRMS), where episodes of neurologic dysfunction are separated by periods of remission (Baecher-Allan et al., 2018). Within a decade of first diagnosis, the majority of RRMS patients develop progression of clinical deficits (Coret et al., 2018; Zhang et al., 2015). The progressive form of MS is refractory to approved RRMS disease-modifying therapies and mechanisms of clinical progression in MS are poorly understood (Goldschmidt and McGinley, 2021).

CD11c^+^ microglia are unique microglia that emerge in the CNS during development and injury (Butovsky et al., 2006; Hagemeyer et al., 2017; Kamphuis et al., 2016; Wlodarczyk et al., 2015; Wlodarczyk et al., 2017). CD11c^+^ microglia have poor antigen presenting capacity (Ford et al., 1995; Giles et al., 2018; Greter et al., 2005; Miller et al., 2007) and increase in CD11c^+^ microglia precedes the onset of experimental autoimmune encephalomyelitis (EAE) an animal model of MS (Fischer and Reichmann, 2001; Serafini et al., 2000). A growing number of evidences suggests that CD11c^+^ microglia possess a neuroprotective function in disease. Increase in CD11c^+^ microglia reduced clinical disability in EAE (Wlodarczyk et al., 2018) and alleviated cuprizone-induced demyelination (Sato-Hashimoto et al., 2019). CD11c^+^ microglia were identified on site of amyloid plaques preventing plaque formation in a transgenic model of Alzheimer’s disease (Butovsky et al., 2006; Kamphuis et al., 2016; Keren-Shaul et al., 2017). The function of CD11c^+^ microglia in autoimmune demyelination is not well understood.

Among the key regulators of inflammation is mitogen-activated protein kinase (MAPK) 14 or p38α (Zarubin and Han, 2005). p38α expression is upregulated in active and chronic lesions in patients with MS (Lock et al., 2002). p38α signaling in CD11c^+^ classical dendritic cells (cDC) is required for the differentiation of uncommitted T cells toward T_H_17 phenotype in MS and EAE and involves the IL-6 – STAT3 signaling pathway (Di Mitri et al., 2015; Huang et al., 2012). p38α is also required for the reactivation of myelin-primed T_H_17 cells by cDC in adoptive EAE transfer (Huang et al., 2012). A sex-dependent role of p38α in EAE has been reported in myeloid cells, where pharmacological inhibition or myeloid cell knockout of *p38α* ameliorated EAE clinical symptoms and inflammation in female mice (Krementsov et al., 2014). The role of p38α in EAE/MS progression remains unknown.

Here we report that a subgroup of female mice with knockout of p38α in CD11c^+^ cells (*KO*), initially protected from EAE, develop EAE progression around 30 days post immunization (dpi). We further describe clinical progression in female and male C57BL/6 wild type (*WT*) mice with initially mild EAE. A reduction in CD11c^+^ microglia and dispersed coalescent parenchymal infiltration were evident at EAE progression in both conditions. In animals that did not develop EAE progression, we found CD11c^+^ microglia in association with astrocyte processes and vasculature. Together, our data indicate a protective role of CD11c^+^ microglia in controlling immune cell parenchymal infiltration during chronic autoimmune demyelination.

## Materials and Methods

### Animals

Female and male C57BL/6 *WT* (JAX stock #000664), *p38α*^fl/fl^ (*Floxed*) ([B6.129-*Mapk14*<tm1.2Otsu], Riken #RBRC02192), and *p38α*^ΔCD11c^ (*KO*) mice 12 weeks old were used to study EAE progression. Mice carrying the *p38α floxed* allele were crossed with mice expressing Cre recombinase under the CD11c promoter ([C57BL/6J-Tg(*Itgax*-cre-EGFP)4097Ach/J], JAX stock #007567). Offsprings were genotyped for the homologous presence of *p38α floxed* allele and *Cre* insert as described earlier (Lo et al., 2014). Animals were maintained in accordance with the NIH Guide for the Care and Use of Laboratory Animals. Experimental protocols were approved by the Institutional Animal Care and Use Committee at the University of California, Davis.

### EAE induction

EAE was induced in female and male mice at the age of 12 weeks as described earlier with some modifications (Chechneva et al., 2011). In brief, mice were injected subcutaneously at two flank sites with an emulsion of myelin oligodendrocyte glycoprotein peptide fragment that spans amino acids 35-55 (MOG) (New England Peptide) in incomplete Freund’s adjuvant supplemented with 500 µg Mycobacterium tuberculosis H37Ra (Difco). Our previously described EAE model with MOG prepared by sonication causes severe EAE in C57BL/6 mice (Chechneva et al., 2011). Sonication changes biochemical composition of MOG epitopes and/or Mycobacterium tuberculosis causing an enhanced immune response (Adelmann et al., 1995). In *KO* mice, sonicated MOG causes mild EAE. EAE in *KO* and *Floxed* control mice was induced by MOG (300 μg) emulsion prepared by quick sonication (3×5 sec on ice). To induce mild EAE in *WT* mice, MOG (150 μg) was homogenized with 2 syringes connected by a stopcock (twice 10 min with 3 h interval on ice). Emulsion was kept in the dark at 4°C between mixing and before preparing into individual doses in syringes. All mice received 200 ng of pertussis toxin (PTX) (List Biological Labs) intraperitoneally (i. p.) on day 0 and 2. The severity of clinical symptoms was evaluated using following neurologic scoring for EAE: 0 - no deficit; 0.5 - mild loss of tail tone; 1 - complete loss of tail tone; 1.5 - mildly impaired righting reflex; 2 - abnormal gait (ataxia) and/or impaired righting reflex; 2.5-single hind limb paresis and ataxia; 3 - double hind limb paresis; 3.5 - single limb paralysis and paresis of second limb / paresis of both hind limbs and weakness of front limbs; 4 - full paralysis of two limbs; 4.5 - hind limb paralysis with hind body paresis; 5 – moribund or death. Animals were monitored and scored daily. At 30 dpi, *WT* mice were organized in three groups, according to clinical phenotype. Mild EAE was defined by chronic mild clinical symptoms throughout EAE course, ranging from loss of tail tone to abnormal gait/paresis (maximum score 3). Severe EAE included animals with minimal recovery from EAE peak and scores of 3.5 (single limb paralysis) and above throughout chronic EAE course. EAE progression was identified by transition from mild to severe disease around 30 dpi with an acute increase in clinical score by 1-2.5 points.

### Histology and pathological scoring

Histological analysis was performed as described earlier (Chechneva et al., 2011). Briefly, mice were anesthetized with isoflurane and perfused with ice cold phosphate buffered saline (PBS). Lumbar spinal cords were dissected, cryopreserved in 20% sucrose, embedded in OCT compound and cut in 15 µm sections on the cryostat. For Luxol Fast Blue (LFB) staining, sections were incubated with 0.1% LFB solution at 60°C for 16 h followed by differentiation with 0.05% lithium carbonate and co-stained with 0.1% Cresyl Violet (Nissl). Slides were mounted using Cytoseal-60 medium and images were acquired on Keyence BZ-9000 microscope. For pathological assessment, for each animal, six sections of the lumbar spinal cord in the interval of 180 µm were scored by blinded investigator for infiltration, vacuolation caused by swollen axon sheaths and demyelination. We used following scoring system: 0 - absence of lesion and infiltration; 1 - not clearly visible lesions and infiltration, 2 – clearly visible lesions and infiltration, 3 – prominent multifocal lesions and infiltration, 4 – marked continuous lesions and infiltration (Anderson et al., 2011).

### Immunohistochemistry

For immunostaining, sections were washed with PBS to remove residues of OCT compound. Nonspecific binding of antibodies was blocked using 10% normal serum in 0.2% Triton-X in PBS. Samples were incubated with rabbit anti-Iba1 (ionized calcium binding adaptor molecule 1; 1:500; Wako 019-19741), rabbit anti-CD4 (1:200; Abcam ab183685), rat anti-CD4 (1:100; BD Pharmingen 550280), rat anti-CD31 (1:200; Abcam ab56299), chicken anti-GFP (1:500, Abcam ab13970), rabbit anti-CCL2 (1:200, Millipore AB1834P), DyLight 594-Lectin (1:200, Vector Laboratories DL-1177) overnight at 4 °C. Fluorophore- conjugated secondary antibodies Alexa Fluor 488, 555 or 647 (1:1000; Life Technologies) were used to detect primary antibody. Slides were mounted using DAPI Fluoromount-G (SouthernBiotech). Images were acquired on Nikon Eclipse 90i (A1) microscope. For DAB staining, sections were treated with 1% H_2_O_2_ to stop endogenous peroxidase activity before incubation with blocking serum. Samples were incubated with primary antibody. HRP- conjugated secondary antibody was used to detect primary antibody. Positive signal was visualized using ImmPACT DAB Peroxidase substrate kit (Vector labs). Slides were mounted using Cytoseal-60 medium and images were acquired on Keyence BZ-9000 microscope. For fluorescent and DAB staining negative controls omitting primary antibody were performed. Images were processed using Adobe Photoshop CC.

### Flow cytometry analysis

Mice were deeply anesthetized and transcardially perfused with ice-cold PBS for 3 min with a flow rate at 3 ml/min. All steps were performed on ice (4°C), unless otherwise stated. Brain and spinal cord were dissected and placed in ice-cold HBSS (Hank’s Balanced Salt Solution) in 2.5 cm culture dish. For tissue digestion, papain resulted in a higher cell yield compared to collagenase. However, we found that papain digestion resulted in the loss of CD11c expression on the cell surface. Therefore, CNS tissue for all FACS experiments were digested using collagenase. Collagenase D (Roche) 1 mg/ml and 0.02 mg/ml with 100 U/ml DNAse I (Worthington) were added to each dish (final concentration), tissue was chopped into ∼1-2 mm^3^ pieces and incubated while shaking at 100 rpm at 37 °C with 5% CO_2_ for 20 min. Enzymatic activity was blocked by EDTA at final concentration 2 mM. Tissue was homogenized in Dounce homogenizer (5-8 strokes) and cell suspension was passed through a 100 µm cell strainer (Corning) in 50 ml falcon tubes. Cells were collected at 350 x g for 5 min at 4°C, resuspended in 37% Percoll (GE Healthcare), transferred to 15 ml falcon tubes and centrifuged at 500 x g for 15 min with 0 brake at 4°C. Cell myelin layer containing cell debris was aspirated. Cell pellet was resuspended in cold 0.5 ml RBC lysis buffer (Biolegend, 420301), transferred to 15 ml falcon tubes and incubated for 30 sec. 5 ml PBS was added before centrifugation at 350 x g at 4°C for 5 min. Cells were resuspended in FACS buffer (PBS plus 1% BSA and 0.1% sodium azide), counted using hemocytometer and transferred to staining plates (Fisher Scientific, 12565502). Following 15 min of Fc receptors blocking with CD16/CD32 antibody (BD Pharmingen), cells were stained with fluorophore-conjugated antibodies: CD11b BV605 (1:200, clone M1/70, Biolegend), CD45 PerCP-Cy5.5 (1:200, clone 30-F11, Biolegend), Ly6G APC-Cy7 (1:200, clone 1A8, BD Biosciences), CD11c FITC (1:100, clone N418, Biolegend), CD11c APC (1:100, clone N418, Biolegend) for 30-60 min. Cells were washed twice with FACS buffer and once with PBS to remove residues of BSA, then incubated with dead cell dye, Zombie red (1:1000 in PBS, Biolegend) for 15 min. After washing, cells were analyzed on Attune NxT Acoustic Focusing Cytometer (Life Technologies). For cytokine detection, cells were counted and incubated in RPMI-1640 medium supplemented with 10% FBS, 2 mM glutamax, 1 mM sodium pyruvate, 0.1 mM NEAA, 100 U penicillin/streptomycin, 50 µM β-ME with Golgi plug (BD Biosciences, 555029) for 1 h at 37^0^C with 5% CO_2_. Phorbol 12-myristate 13-acetate (PMA, 50 ng/ml final concentration) and ionomycin (750 ng/ml final concentration) were added to incubate for 4-5 h at 37°C. Fc receptors were blocked as described above and cells were stained with CD3e FITC (1:200, clone 145-2C11, BD Pharmingen) or CD3e PE-Cy7 (1:200, clone 145-2C11, BD Pharmingen). Dead cells were labeled using Violet fluorescence reactive dye for 20 min (1:1000, ThermoFisher, Cat# L34963). Cells were fixed for 15 min using BD Cytofix/Cytoperm reagent (BD Biosciences, 554722). After washing with BD Perm/Wash (BD Biosciences, 554723) twice, cells were incubated overnight at 4°C with IFN-γ APC (1:25, Clone XMG1.2, Biolegends) and IL-17 PE (1:50, Clone TC11-18H10, BD Pharmingen) in Perm/Wash buffer. Cells were washed and resuspended in FACS buffer for analysis on Attune NxT Acoustic Focusing Cytometer (Life Technologies). Unstained cells, single stain controls and stained AbC Total Antibody Compensation beads (Thermo Fisher, A10497) were used for each experiment to determine compensation parameters. For isotype control of fluorescence minus one, antibody was substituted by fluorophore-conjugated isotype control. Acquired data were analyzed using FlowJo software (TreeStar). To define populations of CD11c^+^ microglia, we found gating on CD11b^+^ and CD45^+^ cells from the pool of CD11c^+^ cells was more accurate than gating of CD11c^+^ cells from CD11b^+^CD45^low^ due to the lack of clear separation of macrophages/monocytes from microglia within CD11b^+^CD45^int^ population. Absolute numbers of cell types isolated from the CNS was determined by multiplying the total number of live cells to the proportion of the cell type acquired by flow cytometer. Quantitative data were presented in absolute numbers.

### Proliferation assay of CD11c^+^ cells

EdU is a thymidine nucleoside analog that incorporates into DNA during S-phase of the cell cycle. To label proliferating cells, animals were pulsed twice with 12 mg/kg EdU intraperitoneal injection in 12 h interval. 24 h after the first injection, animals were deeply anesthetized and perfused with ice-cold PBS. Brain and spinal cord were dissected, and cells were isolated as described above. Following steps were performed on ice (4°C), unless otherwise stated. Fc receptors were blocked with CD16/CD32 antibody (BD Pharmingen) for 15 min and cells were stained with fluorophore-conjugated antibodies: CD11b BV605 (1:200, clone M1/70, Biolegend), CD45 PerCP-Cy5.5 (1:200, clone 30-F11, Biolegend) and CD11c APC (1:100, clone N418, Biolegend) for 30 min. Zombie violet (1:1000, Biolegend) was used to label dead cells. Cells were then fixed with 4% PFA for 15 min at RT. After washing with 1% BSA in PBS, cells were permeabilized with saponin-based permeabilization and wash reagent (Invitrogen, Click-iT Plus EdU Kit) for 15 min at RT. Residues of permeabilization buffer were removed by centrifugation at 350 x g for 5 min. EdU was detected by incubation with freshly prepared Click-iT cocktail containing Tris-HCl (100 mM, pH 7.6), 2 mM CuSO_4_, 2.5 µM Sulfo-Cy3 Azide (Limiprobe) and 100 mM ascorbic acid for 15 min at RT in the dark. Cells were washed with FACS buffer and analyzed on Attune NxT Acoustic Focusing Cytometer (Life Technologies). For Cy3 single stain compensation control we used cells isolated from bone marrow. Proliferation rate was determined as the percentage of EdU^+^ cells within cell populations.

### p38 kinase activity

p38 MAPK activity was assessed by quantifying phospho-p38 MAPK expression levels. *KO* and *Floxed* female and male mice were intraperitoneally injected with 3 mg/kg LPS (O111:B4, Sigma). 24 h after, mice were deeply anesthetized and spleens were isolated, chopped in ice-cold HBSS, and enzymatically digested for 20 min at 37°C. Digested tissue was passed through a 70 µm cell strainer into 50 ml falcon tubes using the backside of 1 ml syringe. Cells were pelleted by centrifugation at 350 x g at 4°C for 5 min and resuspended in ice-cold 1 ml RBC lysis buffer. After washing and centrifugation, cells were resuspended in MACS buffer (PBS with 0.5% BSA and 2 mM EDTA) and counted using hematocytometer. After centrifugation, cells were resuspended and incubated in MACS buffer containing Fc blocker and CD11c MicroBeads (Miltenyi Biotec) according to manufacturer instruction. CD11c^+^ cells were isolated using LS columns (Miltenyi Biotec, Cat# 130-108-338), pelleted and processed for staining on ice (4°C). After Fc receptor blocking with CD16/CD32 antibody (BD Pharmingen), cells were incubated with fluorophore-conjugated antibodies CD11b BV605 (1:200, clone M1/70, Biolegend) and CD11c APC (1:100, clone N418, Biolegend) for 30-60 min. After washing, cells were incubated with dead cell dye Zombie red (1:1000 in PBS, Biolegend) for 15 min. Cells were fixed for 15 min using BD Cytofix/Cytoperm reagent. After washing with BD Perm/Wash, cells were incubated overnight with phospho-p38 MAPK PE (Thr180/Tyr182 (3D7), 1:50, Cell Signaling). Cells were analyzed on Attune NxT Acoustic Focusing Cytometer (Life Technologies). Unstained cells, single stain controls, stained AbC Total Antibody Compensation beads and isotype controls were used for each experiment. Acquired data were analyzed using FlowJo software (TreeStar).

### RNA isolation and qPCR

RNA transcription levels of CCL2 in the lumbar spinal cord was analyzed using qPCR as described earlier (Chechneva et al., 2011). Briefly, mice were anesthetized and transcardially perfused with ice-cold PBS. Lumbar spinal cords were dissected and stored separately in liquid nitrogen. Total RNA was isolated from spinal cord tissue using Direct-zol RNA MiniPrep Kit (Zymo Research) following the manufacturer protocol. RNA at purity OD_260/280_ ratio between 1.8 and 2.0 was reverse-transcribed to cDNA using Multiscribe reverse transcriptase (Applied Biosystems). TaqMan gene expression assay (Applied Biosystems) was used for *CCL2* (Mm00441242_m1). All experimental samples were normalized to the expression level of *GAPDH* (4352339E Applied Biosystems). Relative quantification of fold-change was performed by applying the 2^-ΔΔ^CT method and comparing Cp values of individual mice (Livak and Schmittgen, 2001).

### Statistics

Statistical analysis was performed using GraphPad Prism 7.5. The numbers of animals per group used for each study are shown in the figures as individual dots or indicated in the figure legend. EAE clinical scores were pooled from four independent EAE experiments and evaluated for statistical significance using Two-way ANOVA with Sidak’s multiple comparisons test. One-way ANOVA with Tukey’s post hoc test was used to analyze data in Table 1. Two group experiments were analyzed using Student’s t-test. Experiments with three and more groups were analyzed using One-way ANOVA with Tukey’s post hoc test. In all figures, data are represented as mean ± s.e.m. In all cases, probability values less than 0.05 were considered statistically significant. Images were assembled using Adobe Illustrator CC.

**Table 1.** Clinical assessment of EAE in *p38α Floxed* and CD11c promoter-driven *p38α KO* female and male mice.

| Group | n | Onset | Mean score 1<sup>st</sup> peak | Mean score d26 | Mean score d32 | Max severity from d29 | Duration of paralysis d29 – d60 | Score at d60 | Survival by d60 |
| --- | --- | --- | --- | --- | --- | --- | --- | --- | --- |
| *Floxed* males | 41 | 11.2 ± 0.2 | 4.2 ± 0.1 | 3.2 ± 0.2 | 3.4 ± 0.2 | 3.9 ± 0.1 | 25.9 ± 1.8 | 3.7 ± 0.2 | 90.2% |
| *Floxed* females | 32 | 12.4 ± 0.4 | 4.0 ± 0.1 | 3.2 ± 0.2 | 3.3 ± 0.2 | 3.7 ± 0.2 | 23.9 ± 2.3 | 3.4 ± 0.2 | 90.6% |
| *KO* males | 47 | 12.2 ± 0.3 | 3.3 ± 0.1<sup>***</sup> | 2.2 ± 0.2<sup>***</sup> | 2.1 ± 0.2<sup>***</sup> | 2.7 ± 0.2<sup>***</sup> | 10.0 ± 2.1<sup>***</sup> | 2.1 ± 0.2<sup>***</sup> | 95.7% |
| *KO* females Mild | 17 | 13.2 ± 0.4<sup>*</sup> | 2.9 ± 0.2<sup>***</sup> | 2.0 ± 0.2<sup>***</sup> | 2.1 ± 0.1<sup>**</sup> | 2.3 ± 0.1<sup>***</sup> | 0 <sup>***</sup> | 2.0 ± 0.2<sup>***</sup> | 100% |
| *KO* females Prog | 23 | 14.6 ± 0.8<sup>###</sup> | 3.6 ± 0.2<sup>\$</sup> | 2.6 ± 0.3 | 3.5 ± 0.2<sup>###</sup> | 4.1 ± 0.1<sup>###</sup> | 19.4 ± 2.5<sup>#</sup> | 3.7 ± 0.2<sup>###</sup> | 91.3% |

Data are the mean ± s.e.m. \**P*<0.05, \*\**P*<0.01, \*\*\**P*<0.001, comparing between same sex *Floxed* and *KO* mice. #*P*<0.05, ###*P*<0.001, comparing between females and males of the same genotype. \$*P*<0.05, comparing between *KO* females with and without progression. *P* values were determined by One-way ANOVA with Tukey's post hoc test. d, day after immunization; Prog, progression.

## Results

### Female mice with knockout of *p38α* in CD11c^+^ cells are susceptible to EAE progression

We induced EAE in *KO* and *Floxed* female and male mice and followed EAE clinical symptoms for 60 days. As reported earlier, *Floxed* mice developed chronic severe disease, while *KO* mice were protected and initially displayed mild EAE (Huang et al., 2012; Krementsov et al., 2014) (Fig. 1A, Table 1). EAE onset and peak were delayed in *KO* females compared to *KO* males (Fig. 1A, Table 1). At 30 dpi, 58% of *KO* females underwent clinical progression and transition from mild to severe disease with a sudden increase in clinical score by 1-2.5 points (Fig. 1A, Table 1). No progression of EAE clinical symptoms was observed in *KO* males.

**Fig. 1.**
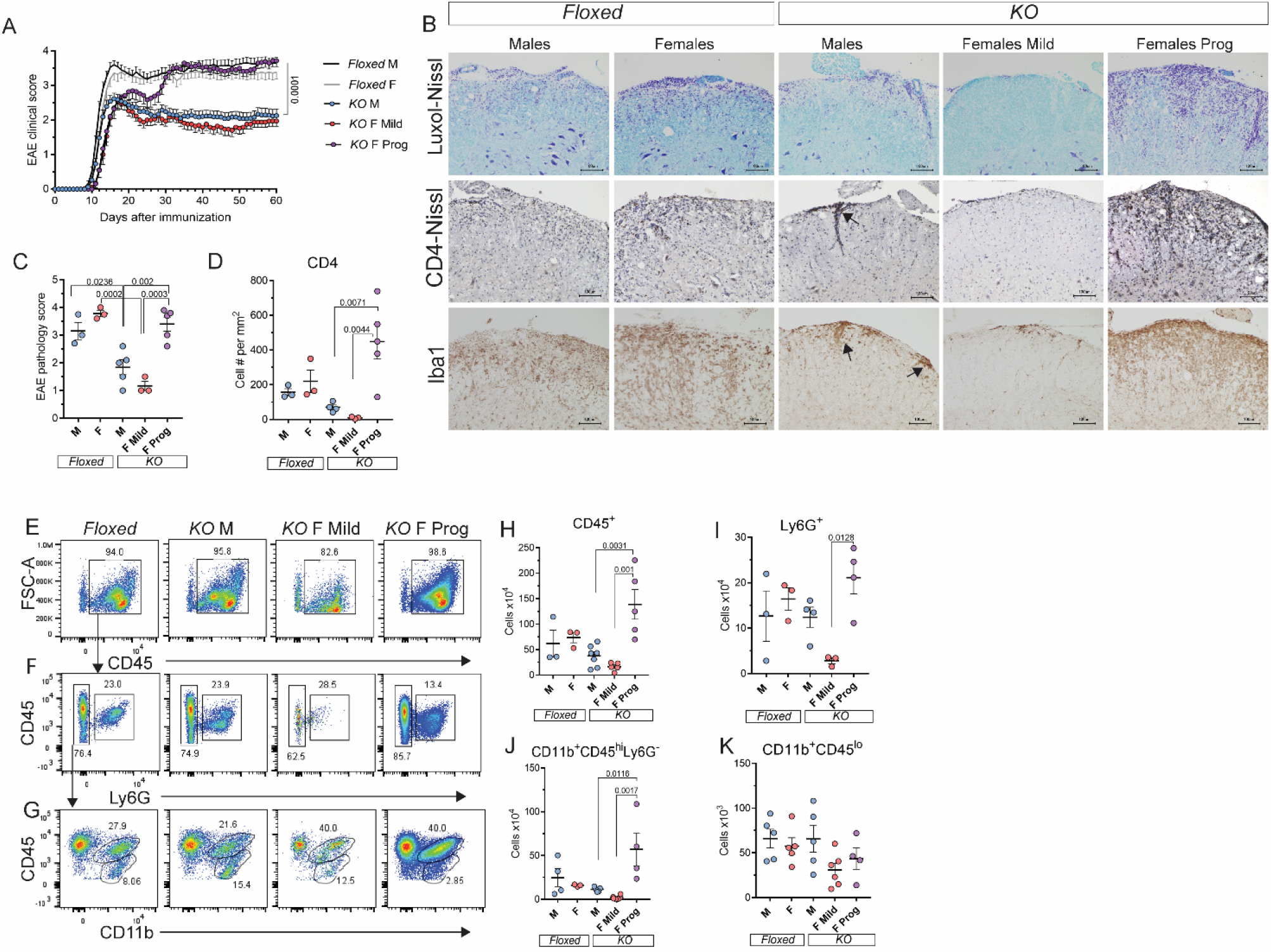
Female mice with knockout of *p38α* in CD11c^+^ cells are susceptible to EAE progression with dispersed parenchymal infiltration. (A) *p38α Floxed* control and CD11c promoter-driven *KO* female and male mice were immunized with MOG prepared by sonication. EAE clinical scores were monitored daily for 60 dpi. Data are representative of four independent experiments with n = 41 for *Floxed* males, n = 32 for *Floxed* females, n = 47 for *KO* males, n = 23 for *KO* females with progression, and n = 17 for *KO* females with chronic mild EAE. *P* values were determined by two-way ANOVA with Sidak’s multiple comparisons test. (B) Representative images of Luxol-Nissl staining, CD4^+^ T cells and Iba1^+^ macrophages/microglia in the lumbar spinal cord at 30 dpi. Clusters of immune cells at the meninges and perivascular areas in *KO* males (arrows). Scale bar 100 µm. (C) Pathological scoring of chronic EAE based on the severity of infiltration and demyelination. Data are representative of two independent experiments. (D) Quantification of CD4^+^ T cells in the white matter parenchyma. (E-G) Flow cytometry analysis of CD45^+^ cells, Ly6G^+^ neutrophils, CD11b^+^CD45^hi^ macrophages/monocytes and CD11b^+^CD45^lo^ microglia isolated from the brain and spinal cord at 30 dpi. (H-K) Quantification of CD45^+^ cells, Ly6G^+^ neutrophils, CD11b^+^CD45^hi^ macrophages/monocytes and CD11b^+^CD45^lo^ microglia. Data are representative of three independent experiments. Each point represents an individual animal. P values were determined by One-way ANOVA with Tukey’s post hoc test. dpi, days post immunization; F, females; M, males; *KO*, knockout; Prog, progression.

We analyzed EAE pathology at 30 dpi in the lumbar spinal cord of *KO* and *Floxed* mice. In progressed *KO* females, we found dispersed parenchymal infiltration, demyelination, and CD4^+^ T cells and Iba1^+^ macrophages/reactive microglia densely scattered throughout the white matter parenchyma (Fig. 1B-D). In contrast, in *KO* males we found a distinct pattern of immune cell infiltration where peripheral immune cells clustered at perivascular areas and the meninges, and CD4^+^ T cells had rare appearance in the parenchyma (Fig. 1B-D). T cells that invaded CNS parenchyma in progressed *KO* females were enriched in INF-γ^+^ T_H_1 population (Supplementary Fig. 1A-D). We found an increase in transcription level of *IL-12b* (also known as *IL-12p40*), a driver of T_H_1 cell differentiation (Balashov et al., 1997) (Supplementary Fig. 1E). In *KO* females with mild EAE that did not transition to EAE progression we observed minor immune cell infiltration (Fig. 1B-D). In *Floxed* mice sustained parenchymal infiltration and demyelination corresponding to chronic severe disease state were evident (Fig. 1B-D).

We further analyzed CNS infiltrates using flow cytometry. CD45^+^ cells were significantly increased in the brain and spinal cord of progressed *KO* females compared to *KO* males and mild non-progressed *KO* females (Fig. 1E, 1H). CD45^+^ cells included Ly6G^+^ neutrophils (Fig. 1F, 1I) and CD11b^+^CD45^hi^ macrophages/monocytes (Fig. 1G, 1J). *KO* females without progression had low numbers of infiltrating immune cells (Fig. 1E-J). There were no significant differences in the numbers of CD11b^+^CD45^lo^ microglia between groups (Fig. 1G, 1K). We conclude that peripheral immune cell infiltration contributes to EAE progression in *KO* female mice.

### Reduction in CD11c^+^ microglia in *KO* female mice

CD11c^+^ cells play an active role in the pathogenesis of MS and EAE (Di Mitri et al., 2015; Giles et al., 2018; Huang et al., 2012; Krementsov et al., 2014; Merad et al., 2013; Mundt et al., 2019). Using flow cytometry, we found an increase in CD11c^+^ cells driven by an increase in the CD11c^+^CD11b^+^CD45^hi^ moDC in the brain and spinal cord of progressed *KO* females (Fig. 2A-D). An increase in the number of CD11c^+^CD11b^-^CD45^hi^ cDC was found in progressed compared to non-progressed *KO* females with mild EAE, reflecting the disease pathology (Fig. 2E). Surprisingly, a significant reduction in CD11c^+^CD11b^+^CD45^lo^ or CD11c^+^ microglia was found in progressed and non-progressed *KO* females (Fig. 2C, F). Gating on CD11c^+^ microglia from the pool of CD11b^+^CD45^lo^ cells resulted in a similar outcome (Supplementary Fig. 2A-B).

**Fig. 2.**
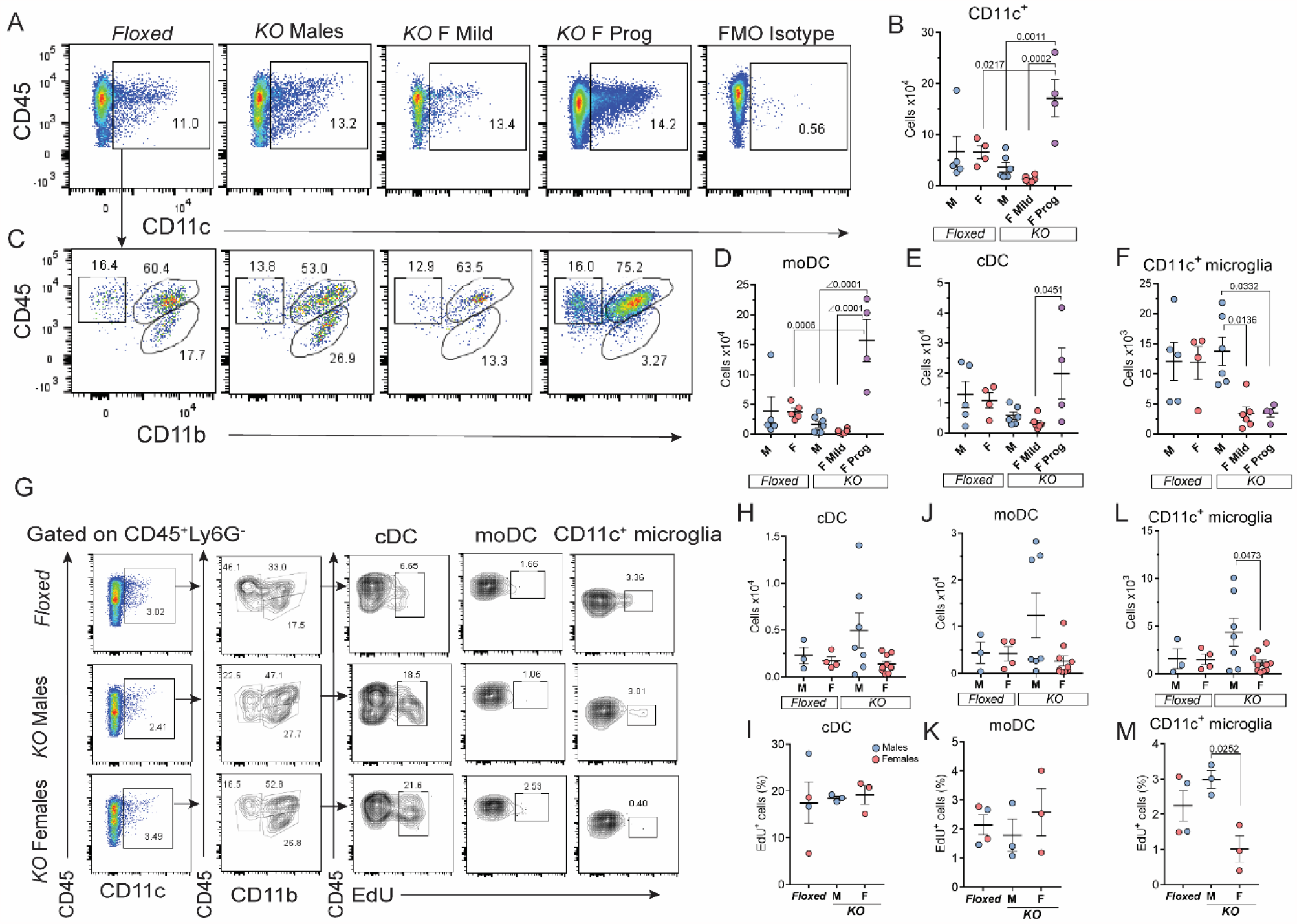
Reduction in CD11c^+^ microglia at EAE progression in *KO* female mice. (A) Flow cytometry analysis of CD11c^+^ cells isolated from the brain and spinal cord of *Floxed* and *KO* male and female mice at 30 dpi. (B) Quantification of CD11c^+^ cells. (C) Flow cytometry analysis of cDC, moDC and CD11c^+^ microglia differentiated by expression of CD45 and CD11b. (D-F) Quantification of moDC, cDC, and CD11c^+^ microglia. (G) MOG-PTX immunized *p38α Floxed* control and CD11c promoter-driven *KO* female and male mice without EAE clinical symptoms at 30 dpi were injected with 12 mg/kg EdU twice in 12 h interval. EdU^+^ cells were analyzed 24 h after first injection. Proliferative response was determined as percentage of EdU^+^ cells of cell population. (H) Quantification of cDC. (I) Proliferative response of cDC. (J) Quantification of moDC. (K) Proliferative response of moDC. (L) Quantification of CD11c^+^ microglia. (M) Proliferative response of CD11c^+^ microglia. Data are representative of three independent experiments on immune cell quantification and two independent experiments on proliferative response. Each point represents an individual animal. *P* values were determined by One-way ANOVA with Tukey’s post hoc test. dpi, days post immunization; F, females; *KO*, knockout; M, males; Prog, progression.

In EAE, microglia increase in numbers by proliferation (Ajami et al., 2011; Jordao et al., 2019). To test whether proliferation of CD11c^+^ microglia is dependent on p38 signaling we compared cell proliferation in MOG-PTX immunized *Floxed* and *KO* mice that did not develop clinical symptoms by 30 dpi. Under this condition, we found an increase in numbers of CNS associated CD45^+^ immune cells, indicating a primed immune response (Supplementary Fig. 3A). The numbers and proliferation rates of CNS associated cDC or moDC were not affected by knockout of *p38α* (Fig. 2G-K). However, we found that numbers and proliferation rate of CD11c^+^ microglia were decreased in *KO* female compared to *KO* male mice (Fig. 2G, 2L-M). Proliferation rate of macrophages/monocytes or total microglia were not different between groups (Supplementary Fig. 3B-D). To demonstrate efficiency of *p38α* knockout, expression of the activated form of p38 MAPK, phospho-p38 MAPK (p-p38), was examined in splenocytes after LPS stimulation. Reduced expression of p-p38 was found in CD11c^+^ splenocytes in *KO* female and *KO* male mice compared to *Floxed* control (Supplementary Fig. 3E-F). We conclude that EAE progression in *KO* female mice is associated with p38α-dependent reduction in CD11c^+^ microglia.

### Reduction in CD11c^+^ microglia occurs at EAE progression in *WT* mice

To study EAE progression in *WT* mice, we induced EAE in C57BL/6 mice with MOG prepared by syringe emulsification. This EAE model resulted in three distinct EAE phenotypes: mild (42%), severe (31%) and initially mild with progression around 30 dpi (27%) (Fig. 3A, Table 2). We compared the pathology between different EAE phenotypes at 30 dpi and EAE peak. Histological analysis revealed dispersed coalescent infiltration of the spinal cord white matter at EAE progression compared to focal infiltrates found at EAE peak (Fig. 3B). A diffuse scattering of immune cells in the parenchyma with lower density than at EAE progression was observed in chronic severe EAE (Fig. 3B). In all EAE groups the vast majority of parenchymal immune cells were Iba1^+^ macrophages/reactive microglia (Fig. 3C). EAE progression was associated with an increase in T_H_1 and T_H_17 cells compared to mild and severe EAE (Supplementary Fig. 4A-D). Flow cytometry showed a significant increase in CD45^+^ cells including neutrophils, macrophages/monocytes, cDC and moDC at EAE peak compared to all chronic EAE groups (Fig. 4A-K). Increase in neutrophils, macrophages/monocytes, cDC and moDC was evident at EAE progression compared to mild and severe EAE (Fig. 4A-K). The numbers of CD11b^+^CD45^lo^ microglia were consistently increased in all EAE groups compared to healthy control (Fig. 4C, 4L). Notable, EAE progression was associated with a significant reduction in CD11c^+^ microglia (Fig. 4E, 4M, Supplementary Fig. 4E-F). There were no sex-dependent differences within immune cell populations between experimental conditions in *WT* mice. We conclude that clinical progression during chronic EAE is associated with dispersed parenchymal infiltration and involves reduction in CD11c^+^ microglia.

**Fig. 3.**
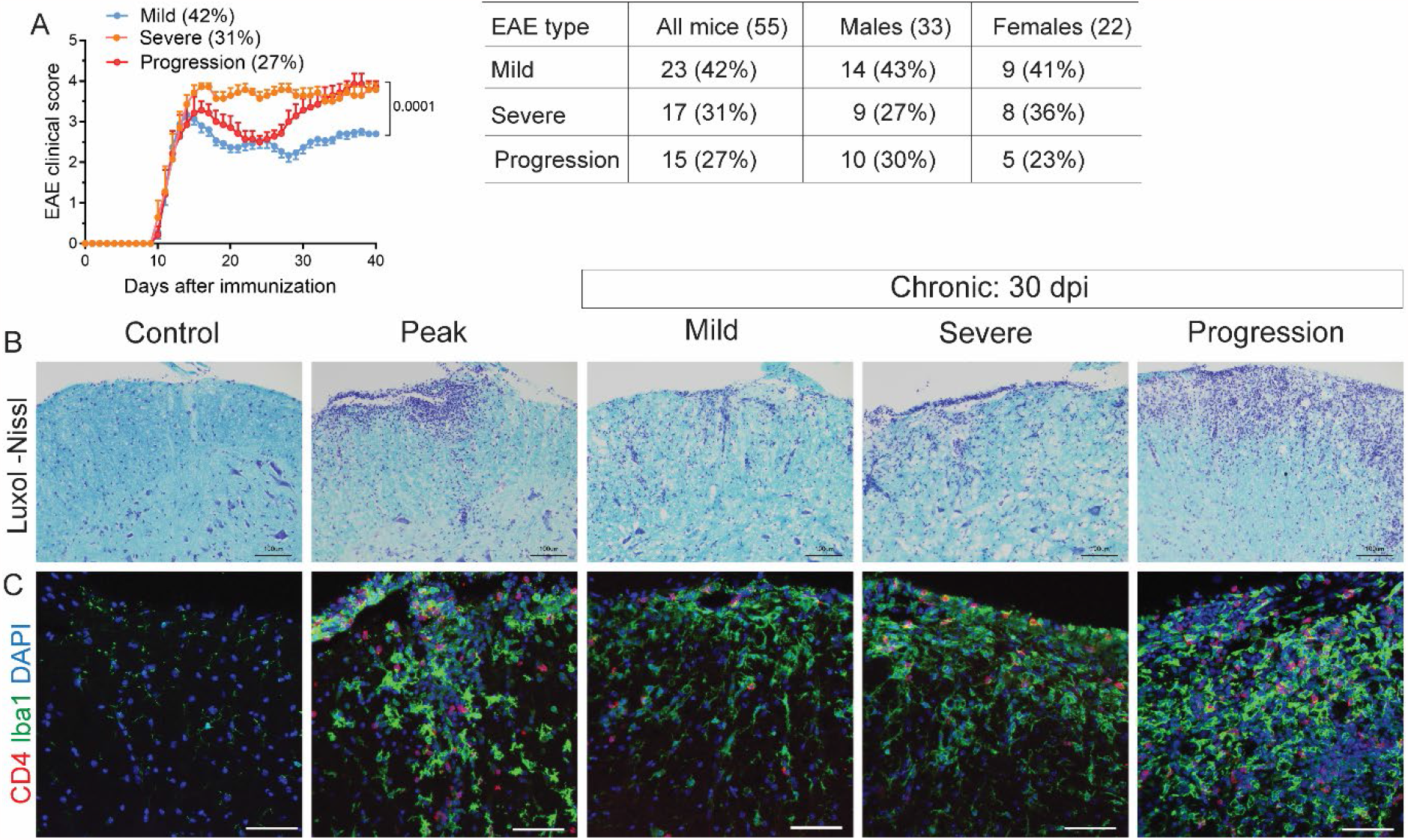
Diffuse parenchymal infiltration at EAE progression in *WT* mice. EAE was induced in C57BL/6 mice with MOG prepared by syringe homogenization. (A) At 30 dpi animals were grouped based on EAE course: mild, severe or progression. Data are representative of six independent experiments. Individual time points represent mean ± s.e.m. *P* values were determined by two-way ANOVA with Sidak’s multiple comparisons test. (B) Representative images showing demyelination and infiltration in the lumbar spinal cord of healthy control, at EAE peak, and in chronic mild, severe and progressive EAE. (C) CD4^+^ T cells and Iba1^+^ macrophages/microglia in the lumbar spinal cord of healthy control, at EAE peak, and in chronic mild, severe and progressive EAE. Scale bar 100 µm for Luxol-Nissl and 50 µm for CD4/Iba1. dpi, days post immunization.

**Table 2.**
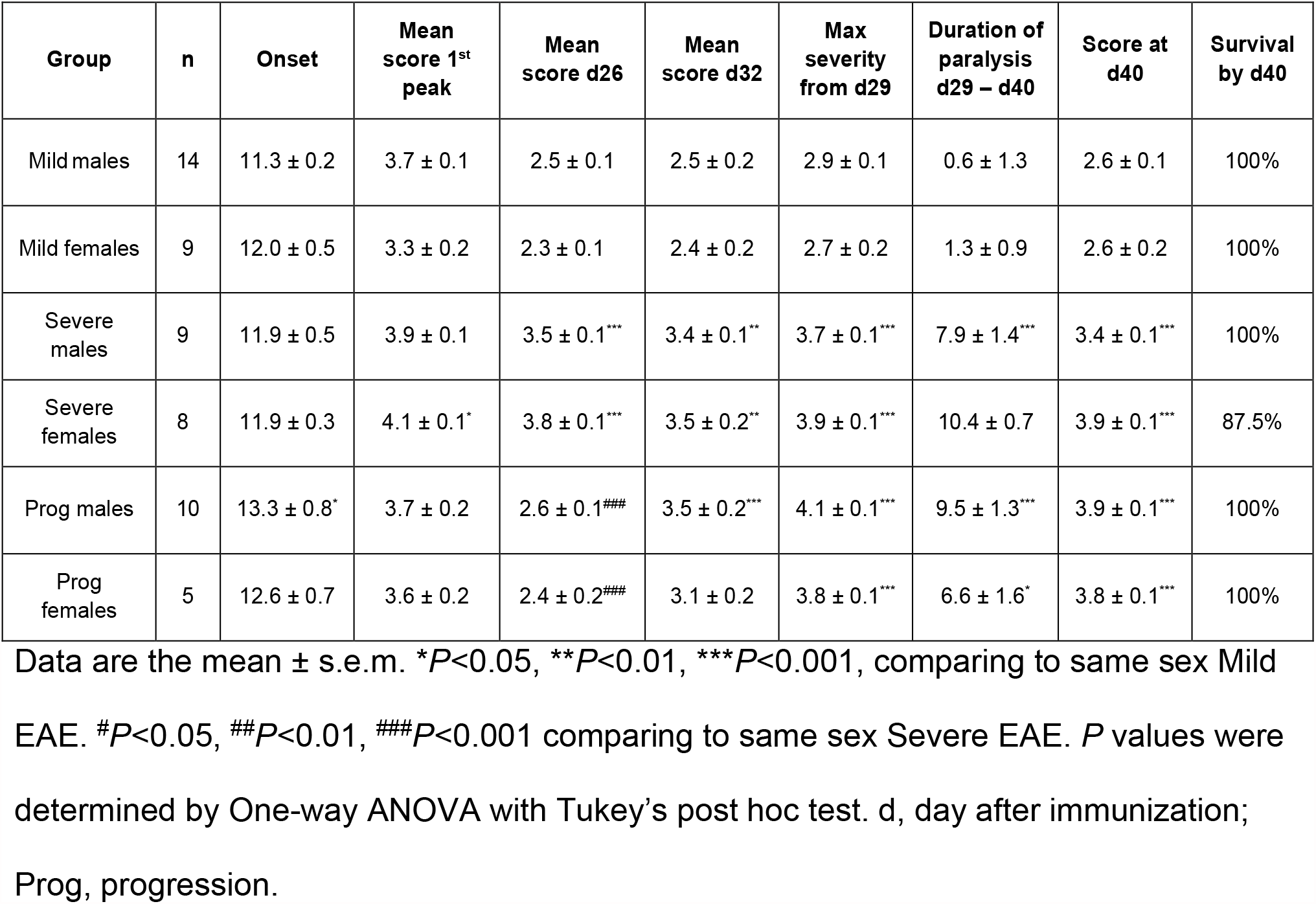
Clinical assessment of EAE in C57BL/6 *WT* mice.

**Fig. 4.**
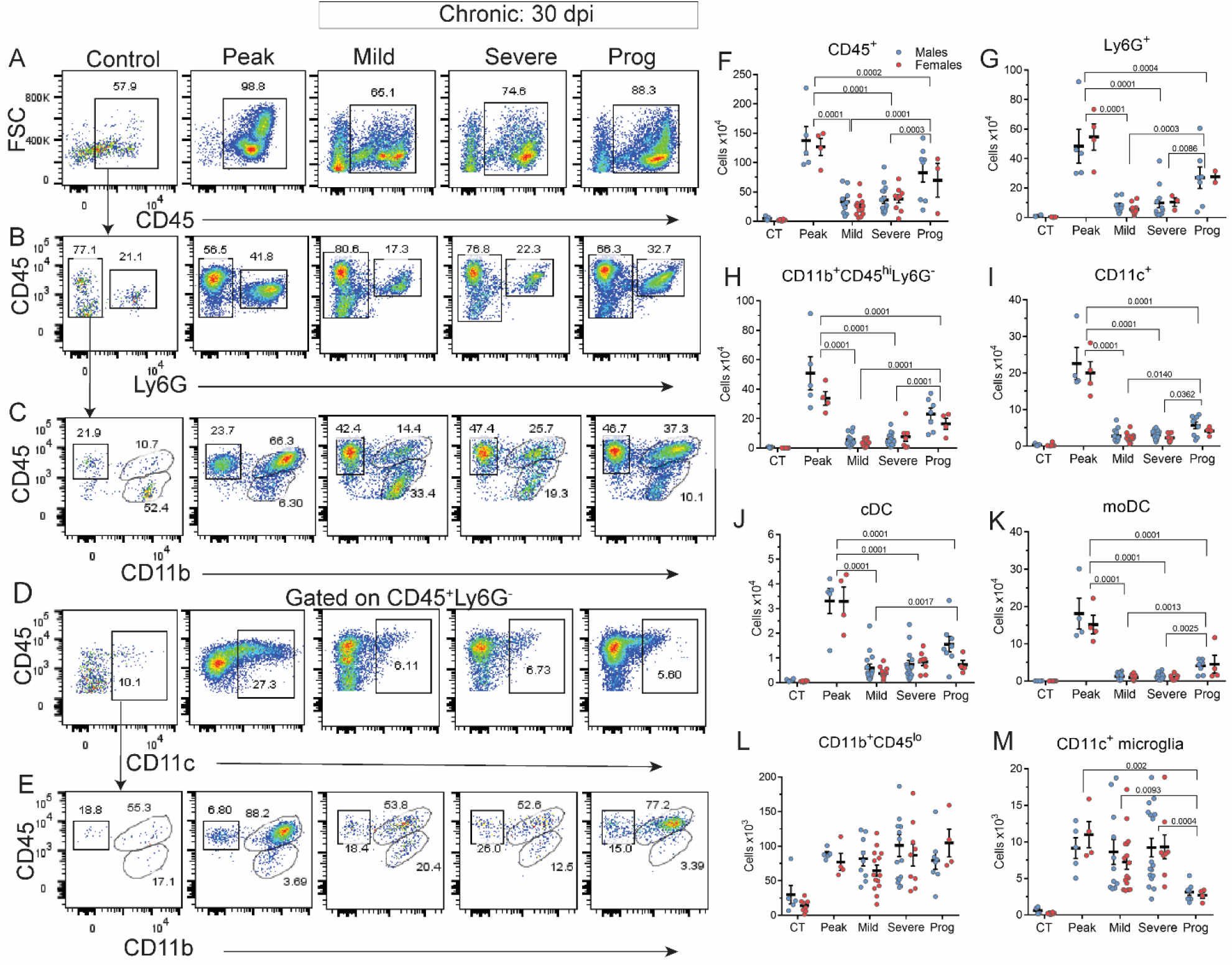
Reduction in CD11c^+^ microglia at EAE progression in *WT* mice. EAE was induced in C57BL/6 *WT* mice with MOG prepared by syringe homogenization. Flow cytometry analysis of cells isolated from the brain and spinal cord of healthy mice, at EAE peak, EAE progression and at 30 days of mild and severe EAE. (A-D) Flow cytometry analysis of CD45^+^ cells, Ly6G^+^ neutrophils, CD11b^+^CD45^hi^Ly6G^-^ macrophages/monocytes and CD11b^+^CD45^lo^ microglia, CD11c^+^ cells. (E) Gating strategy to differentiate cDC, moDC and CD11c^+^ microglia by expression of CD45 and CD11b. (F-M) Quantification of CD45^+^ cells, Ly6G^+^ neutrophils, CD11b^+^CD45^hi^ macrophages/monocytes, CD11c^+^ cells, cDC, moDC, CD11b^+^CD45^lo^ microglia, and CD11c^+^ microglia. Data are representative of six independent experiments. Each point represents an individual animal. *P* values were determined by One-way ANOVA with Tukey’s post hoc test. dpi, days post immunization.

### CD11c^+^ microglia contact astrocyte processes and glia limitans

Astrocytes orchestrate immune cell parenchymal infiltration (Horng et al., 2017; Kim et al., 2014). We examined localization of CD11c^+^ microglia in relation to astrocytes. Commercially available antibodies were not able to detect CD11c^+^ cells by immunohistochemistry, likely due to low expression of CD11c in migrating and activated CD11c^+^ cells (Merad et al., 2013; Prodinger et al., 2011). To visualize CD11c^+^ cells, we used expression of eGFP under the CD11c promoter. In protected *KO* males, we found dendritic-like CD11c^+^ microglia in the vicinity to astrocytes and blood vessels (Fig. 5A-B). CD11c^+^ microglia were abundant in *WT* mice with chronic mild EAE (Fig. 5C). Dendritic-like CD11c^+^ microglia with stellate thick processes formed contacts with astrocytes (Fig. 5D). In progressed *KO* females, disorganized processes of hypertrophic astrocytes were surrounded by infiltrating moDC / ameboid CD11c^+^ microglia (Fig. 5E-F). Dendritic-like CD11c^+^ microglia could not be differentiated from infiltrating moDC by morphology. Infiltration of immune cells into the CNS parenchyma is dependent on astrocyte-derived CCL2 (Dogan et al., 2008; Kim et al., 2014; Moreno et al., 2014). We found upregulation of CCL2 expression in hypertrophic reactive astrocytes in progressed *KO* females (Fig. 5G-H). Our data suggest that CD11c^+^ microglia interaction with astrocytes at the glia limitans is dysregulated in EAE progression.

**Fig. 5.**
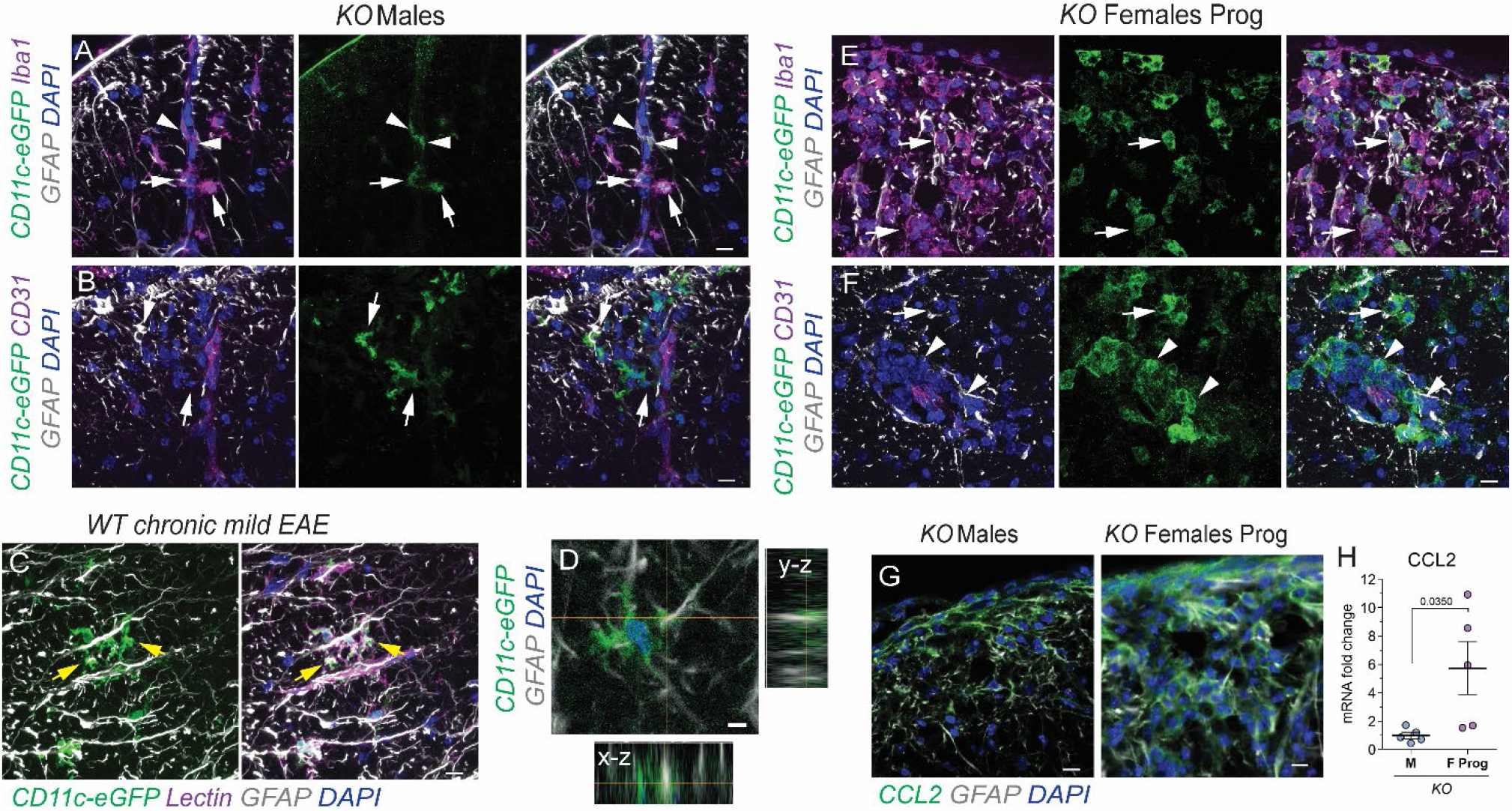
CD11c^+^ microglia form contacts with astrocytes at the glia limitans. CD11c^+^ cells were visualized using CD11c promoter-driven expression of eGFP. (A-B) In the spinal cord white matter of protected *KO* males, parenchymal and perivascular astrocytes were surrounded by CD11c-eGFP^+^ microglia (arrows). moDC (arrowheads) were trapped within perivascular spaces. (C) CD11c^+^ microglia in contact with parenchymal astrocyte processes and glia limitans covering Lectin^+^ blood vessel (arrows) in *WT Itgax*-cre-EGFP mice in chronic mild EAE. (D) Orthogonal projection shows that dendritic-like CD11c^+^ microglia form contacts with astrocyte processes. (E) CD11c-eGFP^+^ moDC/reactive microglia surround hypertrophic processes of reactive astrocytes in the ventral spinal cord of progressed *KO* female mice (arrows). (F) moDC in the perivascular space (arrowheads) and parenchyma (arrows). (G) Increased CCL2 expression in reactive astrocytes in progressed *KO* females. Scale bars 10 µm for A-B, D-G and 5 µm for C. (G) mRNA transcription levels of CCL2 in the lumbar spinal cords of *KO* male and progressed *KO* female mice. Data are representative of three independent experiments. Each point represents one individual animal. P values were determined by Student’s t-test. F, females; *KO*, knockout; M, males; Prog, progression.

## Discussion

EAE allows to study a variety of clinical features of MS (Constantinescu et al., 2011). Here we demonstrate that mice that develop mild EAE after MOG immunization are primed to undergo spontaneous EAE progression around 30 dpi. EAE progression appeared in 58% of female mice lacking *p38α* in CD11c^+^ cells and ∼40% of *WT* mice both of which initially developed mild EAE. Similar progression of clinical symptoms around 30 dpi was reported in non-obese diabetic mice used as a model of EAE progression (Baker et al., 2019; Basso et al., 2008; Buonvicino et al., 2019; Encinas et al., 1999; Tanabe et al., 2019). Why EAE progression follows a 30 dpi pattern and what triggers progression in a fraction of animals with mild EAE remains to be investigated.

Secondary antigenic epitope spreading has been identified as a molecular trigger in MS relapse and progression involving Th1 and Th17 cells (Babaloo et al., 2015; Balashov et al., 1997; McRae et al., 1995; Romme Christensen et al., 2013). Activation of epitope-specific Th cells is dependent on antigen-presenting CD11c^+^ cells (Chastain et al., 2011; Giles et al., 2018; Greter et al., 2005; Mundt et al., 2019) and requires p38α for induction of the Th17 population (Di Mitri et al., 2015; Huang et al., 2012). We found that both, Th1 and Th17 cells were involved in chronic EAE progression in *WT* mice. However, re-activation of Th1 cells through p38α-independent mechanism was enough to trigger EAE progression in *KO* female mice, suggesting mechanistic differences in immune response between initial EAE peak and progression of chronic EAE. *KO* males were protected from EAE progression. Whether Th17 cells are required for EAE progression in males remains to be examined.

Comparing compositions of immune cells and patterns of parenchymal infiltration between EAE phenotypes, we found selective reduction in CD11c^+^ microglia and dispersed coalescent parenchymal infiltration as characteristic features of chronic EAE progression. Importantly, dispersed infiltration but with lesser cell density was also evident in chronic severe EAE, suggesting that dispersed, in contrast to focal infiltration is a pathological hallmark of irreversible disability.

Signaling via p38 MAPK is involved in hormone-dependent sexual dimorphism of the immune response with variable outcomes depending on cell type and condition. In microglia, p38 MAPK was identified as a cause of neuropathic and inflammatory pain in males in chronic constriction injury and mechanical allodynia (Sorge et al., 2015; Taves et al., 2016). Testosterone-dependent increase in p38 MAPK phosphorylation in splenic and peritoneal macrophages was reported in male mice after trauma-hemorrhage (Angele et al., 2003). We found that p38α was involved in maintaining CD11c^+^ microglia in female mice and reduction in CD11c^+^ microglia correlated with increased rate of EAE progression in *KO* females. In contrast, protection from EAE progression in *KO* males was associated with the presence of CD11c^+^ microglia in close contact with astrocytes and at the glia limitans. A biased role of p38α in driving CD11c^+^ microglia reduction in males would explain the lack of EAE progression in *KO* male mice. Defining the molecular mechanism of CD11c^+^ microglia reduction is needed to further understand the sex-specific role of p38α at EAE progression.

In our study, *KO* female mice showed a delay in EAE onset and peak, indicating a delay in the peripheral immune response involving cDC and/or moDC. This is supported by an earlier study where female mice lacking p38α in myeloid cells showed improvement of EAE clinical symptoms while their male cohorts developed exacerbated EAE (Krementsov et al., 2014). Reduced peripheral immune response may explain the lack of EAE progression in the fraction of *KO* females with minor spinal cord infiltration. Changes in the peripheral immune response within *KO* females may be caused by unequal efficiency of *p38α* deletion or compensatory activity of alternative p38 MAPK subunits, such as p38β and p38δ (Huang et al., 2012). Taking together, variable effects of p38α function may explain why many of p38 MAPK inhibitors failed to fulfill their expectations in clinical trials to date and emphasize the need of cell-specific therapeutic approaches (Goldstein et al., 2010; Krementsov et al., 2013).

Dendritic-like CD11c^+^ microglia localized at the glia limitans are found in the brain, spinal cord and optic nerve under healthy conditions and are suggested to play a role in astrocyte glia limitans organization (Prodinger et al., 2011). Glia limitans astrocytic endfeet surround blood-brain barrier endothelial cells and form a physical border between perivascular spaces and CNS parenchyma (Horng et al., 2017). Dendritic-like CD11c^+^ microglia were abundant in the spinal cord of mice with chronic mild EAE, where immune cells were restricted from entering parenchyma. These CD11c^+^ microglia formed physical contacts with astrocyte processes suggesting their direct interaction with astrocytes to limit immune cell parenchymal entry. Expression of triggering receptor expressed on myeloid cells (TREM2) has been associated with CD11c^+^ microglia in EAE and an animal model of Alzheimer’s disease (Keren-Shaul et al., 2017; Piccio et al., 2007). Supporting the concept of a potential protective role of CD11c^+^ microglia in CNS infiltration, diffuse parenchymal infiltration and increase of clinical disability in EAE was found after blockade of TREM2 activity with specific antibody (Piccio et al., 2007).

## Conclusion

Our study provides novel insight into the cellular mechanisms underlying clinical progression during chronic EAE. We identified reduction in CD11c^+^ microglia and the specific pattern of dispersed parenchymal infiltration as pathological hallmarks of EAE progression. Our data suggest a protective role of dendritic-like CD11c^+^ microglia in controlling CNS infiltration and EAE progression through interaction with astrocytes.

## Acknowledgements

This work was in parts supported by grants from National Institutes of Health (R01HD087566 to W.D.) and Shriners Hospital for Children (87310-NCA-19 to O.C.).

## Author contributions

O.V.C. and F.M. designed the research, performed experiments, interpreted the data, wrote and edited the manuscript. Z.D., A.Z., D.J.D., and A.M.H. performed experiments. P.B., D.P., A.S., and W.D. interpreted the data and edited the manuscript.

