## Supplementary Figures 1-4 for "Reduction in CD11c^+^ microglia correlates with clinical progression in chronic experimental autoimmune demyelination"

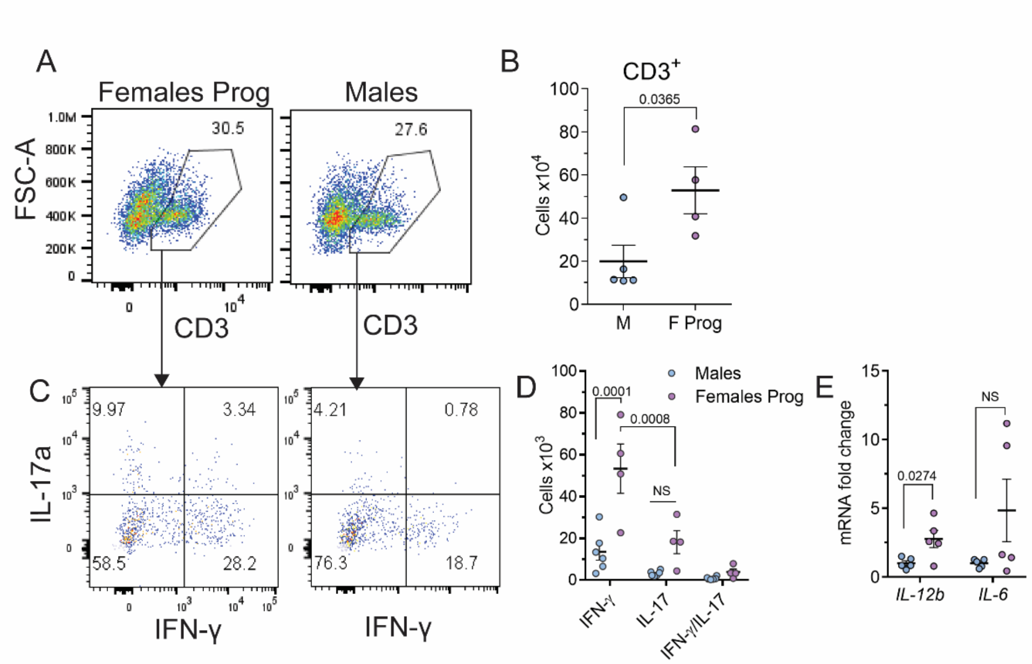


**Supplementary Fig. 1. IFN-$\gamma^+$ cells are increased at EAE progression in KO female mice.**

(A) Flow cytometry analysis of $CD3^+$ T cells in the brain and spinal cord of *KO* male and progressed *KO* female mice. (B) Quantification of $CD3^+$ T cells. (C) Flow cytometry analysis of IL-17$a^+$ and IFN-$\gamma^+$ T cells isolated from the CNS of *KO* male and progressed *KO* female mice. (D) Quantification of IL-17$a^+$ and IFN-$\gamma^+$ T cells. (E) IL-12b and IL-6 mRNA level in the spinal cord of *KO* male and progressed *KO* female mice. Data are representative of two-three independent experiments. *P* values were determined by Student's t-test. Each circle represents an individual mouse and small horizontal line indicates the mean. F, females; M, males; NS, not significant; Prog, progression.

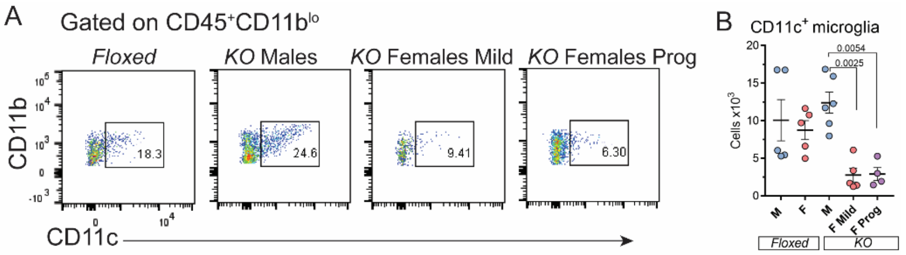


**Supplementary Fig. 2. Reduction in CD11c$^+$ microglia in *KO* female mice.**
FACS analysis of CD11c$^+$ microglia isolated from the brain and spinal cord of *Floxed* and *KO* EAE mice at 30 dpi. (A) CD11c$^+$ cells were gated on CD11b$^+$CD45$^{lo}$ microglia. (B) Quantification of CD11c$^+$ microglia. Data are representative of three independent experiments. Each point represents an individual animal. P values were determined by One-way ANOVA with Tukey's post hoc test. dpi, days post immunization; F, females; *KO*, knockout; M, males; Prog, progression.

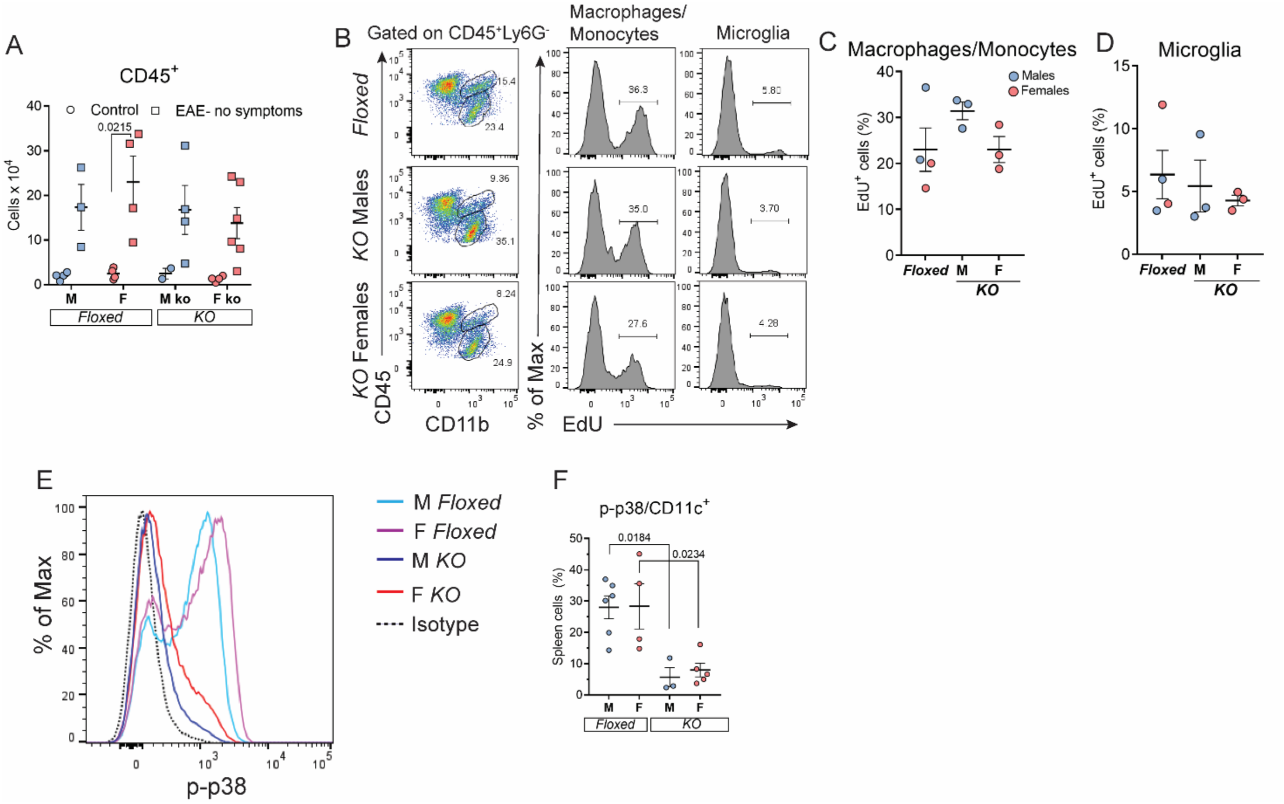


**Supplementary Fig. 3. Proliferation of CNS immune cells during immune response.** Healthy mice and mice without EAE symptoms at 30 dpi were injected with 12 mg/kg EdU twice at 12 h interval. EdU+ cells were analyzed 24 h after first injection. (A) Number of CD45+ cells isolated from the CNS of healthy mice and mice without EAE symptoms at 30 dpi. (B) FACS analysis of EdU+ macrophages/monocytes and microglia in the brain and spinal cord of MOG-immunized mice without EAE symptoms at 30 dpi. (C-D) Proliferation rate of macrophage/monocyte and microglia in mice without EAE symptoms at 30 dpi. (E) *Floxed* and *KO* female and male mice were intraperitoneally injected with 3 mg/kg LPS 24 h before analysis. Expression levels of the activated form of p38 MAPK, phospho-p38 MAPK (p-p38), was examined in CD11c+ splenocytes by flow cytometry. (F) Proportion of p-p38+ CD11c+ splenocytes of total spleen cells. Data are representative of two independent experiments. Each point represents an individual animal. *P* values were determined by One-way ANOVA with Tukey's post hoc test. dpi, days post immunization; F, females; *KO*, knockout; M, males.

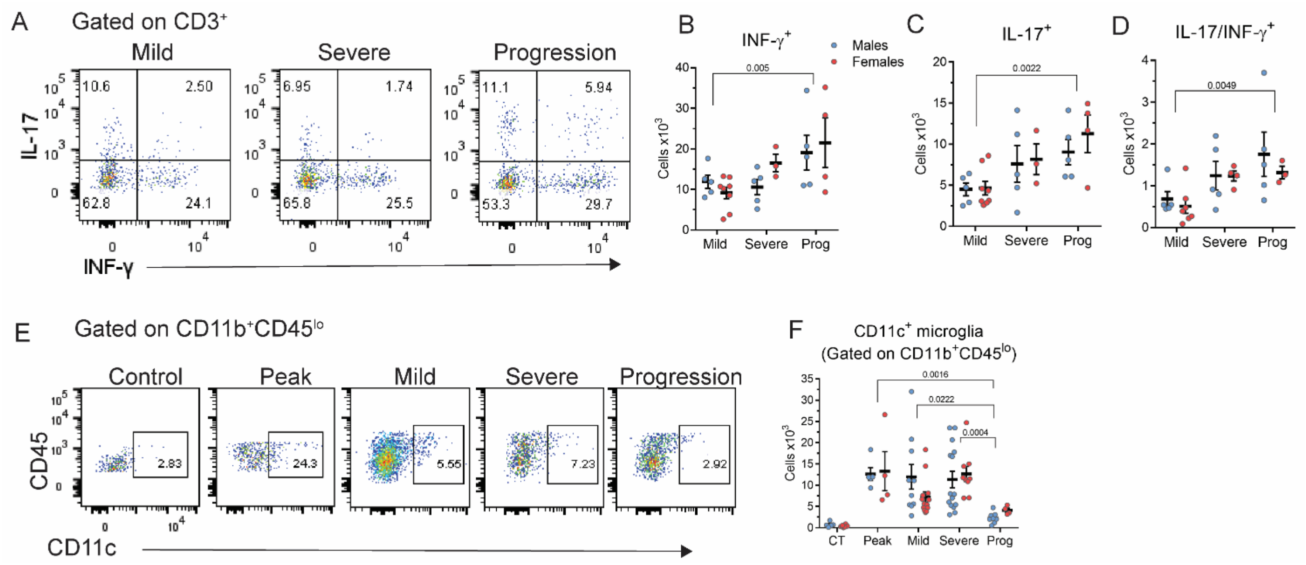


**Supplementary Fig. 4. Increase in T cells and reduction in CD11c+ microglia at EAE progression in *WT* mice.**
(A) Flow cytometry analysis of T cells in the brain and spinal cord of *WT* mice in mild, severe and progressive EAE at 30 dpi. (B-D) Quantification of IFN-γ+, IL-17+ and IL-17/IFN-γ+ T cells. (E) Flow cytometry analysis of CD11c+ microglia gated on CD11b+CD45lo microglia isolated from brain and spinal cord of healthy mice, at EAE peak, and chronic mild, severe and progressive EAE. (F) Quantification of CD11c+ microglia. Data are representative of three (for T cells) and six (for CD11c+ microglia) independent experiments. Each point represents an individual animal. *P* values were determined by One-way ANOVA with Tukey's post hoc test. dpi, days post immunization; Prog, progression.
